# Social tolerance in *Octopus laqueus* - a maximal entropy model

**DOI:** 10.1101/526905

**Authors:** Eric Edsinger, Reuven Pnini, Natsumi Ono, Ryoko Yanagisawa, Kathryn Dever, Jonathan Miller

**Author notes:** These authors contributed equally to this work. EE was lead experimentalist. RP was lead theorist.

## Abstract

*Octopus laqueus* is a small tropical octopus found in Okinawa, Japan and the greater Indo-Pacific. Octopus are often viewed as solitary animals but O. *laqueus* live in close proximity in the wild, and will potentially encounter one another on a regular basis, raising the possibility of sociality in the species. To test for social tolerance and social repulsion in *O. laqueus*, animals were kept in communal tanks, and the number of dens and sex composition was varied per tank, with a set mixture of sizes and with den occupancy tracked per individual. We found that *O. laqueus* will socially tolerate other individuals by sharing tanks and dens, including several animals in contact and sharing a den under den-limited conditions, and with typically no loss to cannibalism or escape. However, animals also exhibit significant levels of social repulsion, and individuals often chose a solitary den when given the option. The patterns of den occupancy are observed to be consistent with a maximum entropy model. Overall, the preference to have a den is stronger than the preference to be solitary in *O. laqueus*, and the animals are socially tolerant of others in the tank and in a den or shelter, a first for octopuses outside mating. The relaxed disposition and social tolerance of O. *laqueus* make it a promising species to work with in lab, and for development into a genetic model for social behavior in octopuses.

## Introduction

Octopus are traditionally viewed as solitary animals that do not form social aggregations, have relatively few and simple reciprocal interactions, and rarely make physical contact outside aggression and mating [1–7]. Further, species are known to be cannibalistic in the laboratory and in the field [8–11]. However, recent studies [12–14] suggest that classifying octopus as merely asocial is overly simplistic. Here and elsewhere in the descriptive sections of this manuscript, we adopt the conventional nomenclature of the field; however, definitions and conventions are not universally shared even within the field, and the predictive value of this nomenclature is unresolved. Our aim is to investigate functional and *predictive* – as opposed to descriptive – characterizations of ‘‘sociality;” such a characterization is proposed in materials and methods, and may not necessarily correspond directly to customary notions of sociality.

‘Asocial’animals by definition reject or lack the capability for social interaction; they are non-interacting, typically ignoring one another. In contrast, some species of octopus show localized aggregated distributions with moderate to high densities, depending on various factors, such as habitat, season, temperature, size, maturity, and prey in the field [1,14–21].

Octopus aggregations are unlikely to represent gregarious attraction between individuals outside mating [1,15,17], but would likely require some amount of social tolerance to minimize frequent aggressive or lethal interactions. Further, the likelihood of individuals running into one another on a daily or frequent basis in a densely localized population is suggestive that more active social interactions, including touch and visual signaling by body color and patterning, do exist. Social tolerance in dense group cultures in lab has also been reported. It was found [22,23], that many animals occupying a single large tank tolerate one another as long as they are well fed and there is not a large size difference between tank mates – a rule of thumb that seems to hold for many octopuses and cephalopods.

Octopuses that are largely solitary and asocial in the field can form dominance hierarchies in the lab. A shift from solitary to hierarchical social structure in the lab suggests that sociality may be a plastic trait in octopuses, one that is flexible or dependent on the conditions at hand, including population density [24–26]. The transition from solitary to hierarchical structure, together with the ability to potentially recognize individuals, can be explained in terms of the “dear enemy phenomenon” [27].

*Octopus laqueus* is a small shallow-water tropical species. It is common in sand, reef rubble, and reef habitats in Okinawa, Japan, and is likely distributed more generally in the tropical Indo-Pacific [2,28]. Animals are common in holes or dens in the sand and reef rubble (Fig 1a) (S1 Video) that were within a few meters or less of one another, suggesting that a given individual is likely to encounter multiple other individuals on a given night of foraging and hunting. The abundance and proximity of *O. laqueus* in the field raised the possibility that it is a social octopus, leading us to ask the specific question, is *O. laqueus* tolerant of other octopus in a den, and to design a series of experiments to test this question. In the following we describe the experiments, develop a statistical model of den occupation based on the maximal entropy principle, and discuss the results.

**Fig 1.**
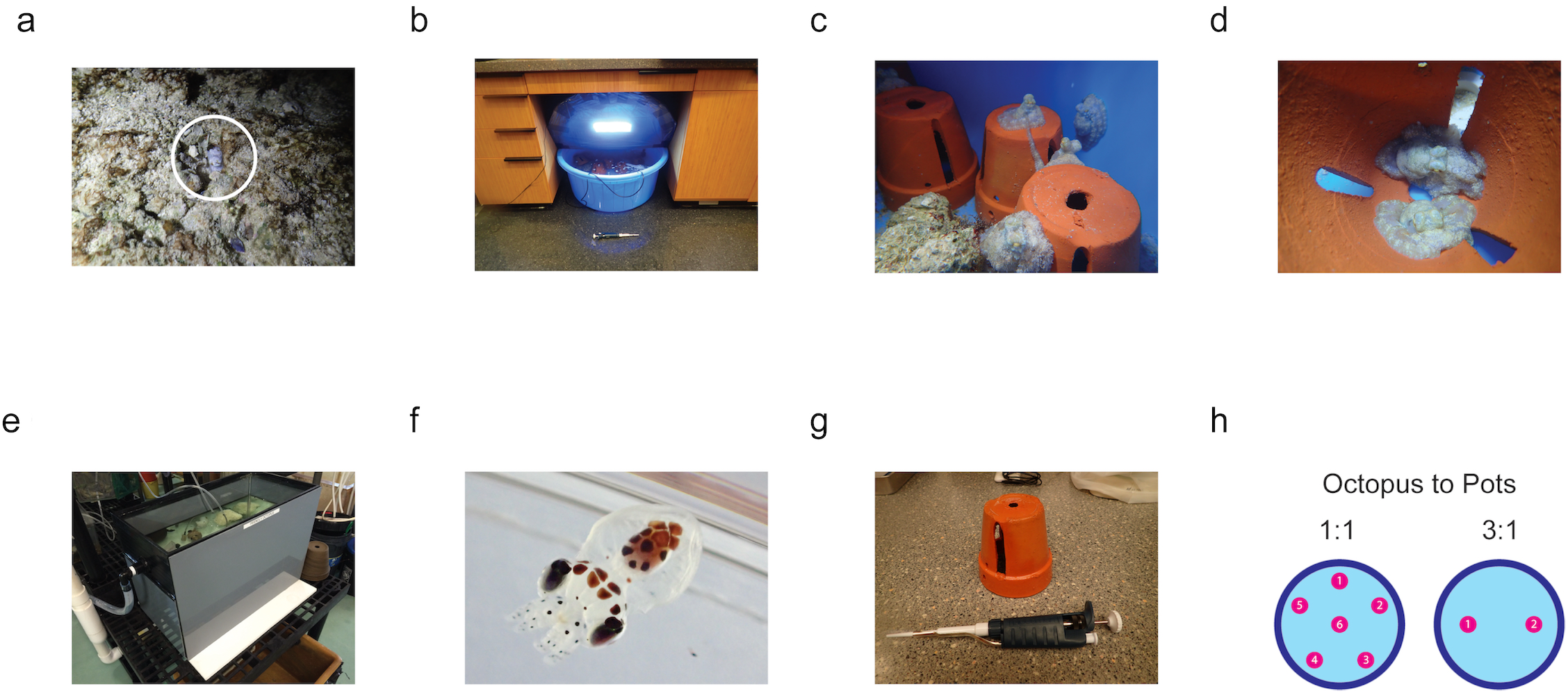
Culturing and tank design of *O. laqueus* in the lab. (a) An adult *O. laqueus* peeking out of its den at Maeda Flats, Okinawa, Japan, where animals were collected for lab culturing. (b) An experimental culturing tank at OIST with air line, tank cover, LED lights, and clay pots visible. (c) Several *O. laqueus* in a tank in lab. (d) Two *O. laqueus* sharing a single pot as a den during the day. (e) The long-term culturing tank at the MBL. (f) An *O. laqueus* hatchling of a female that was ong-term cultured at the MBL, beginning as a juvenile through sexual maturity and ending with a natural death by senescence after hatching of her embryos. (g) A single clay pot with a pipettor for scale. (h) Layout of clay pots in the experimental tanks for the social experiments.

## Methods

### Ethical considerations

The research adhered to ASAB/ABS Guidelines for the Use of Animals in Research, in addition to legal and institutional requirements in Japan and the United States. Collection, care, and export of small non-commercial octopuses, including *O. laqueus*, are not regulated in Japan, and permits or licenses from a granting authority were not required. Import of *O. laqueus* from Okinawa to the Marine Biological Laboratory (MBL) in the United States was done in accordance with all applicable US Customs and US Fish and Wildlife regulations. Care of invertebrates, like *O. laqueus*, does not fall under United States Animal Welfare Act regulation, and are omitted from the PHS-NIH *Guide for the Care and Use of Laboratory Animals.* Thus, an Institutional Animal Care and Use Committee, a Committee on Ethics for Animal Experiments, or other granting authority does not formally review and approve experimental procedures on and care of invertebrate species, like *O. laqueus*, at the MBL. However, in accordance with MBL Institutional Animal Care and Use Committee guidelines for invertebrates, our care and use of *O. laqueus* in Japan and in the United States generally followed tenets prescribed by the Animal Welfare Act, including the three “R’s” (refining, replacing, and reducing unnecessary animal research), and also generally adhered to recent EU regulations and guidelines on the care and use of cephalopods in research [30].

### Collection

*O. laqueus* were collected at night on low tides close to shore in water five to fifty centimeters deep, and in areas of sand and coral rubble in Okinawa, Japan. Collections were made in fall and winter 2014 and November 2015. *O. laqueus* were easily caught when found outside the den, with up to fifty animals collected at a time and placed in a few small buckets with seawater during collection (Fig 1a). Experiments were not initiated until several days after collection and the onset of feeding to allow acclimation to the laboratory environment.

### Culturing

*O. laqueus* were easy to care for in group cultures in lab (Fig 1). Seawater temperature was at room temperature or maintained at 23°C. At the Okinawa Institute of Science and Technology (OIST), animals were kept at densities of up to one animal per 15 liters with four to fifteen animals in 250 liter tanks with filtration, air, and closed circulation. Seawater changes of 10-100% were made every 1-3 days and water quality was checked periodically (pH, nitrates, nitrites, ammonia). Prime (*Seachem*) was periodically used to help stabilize conditions for short-term cultures (days to weeks). For longer-term group cultures of several months, three young juveniles (one male and two females, each around ten grams) were shipped from OIST in Okinawa, Japan to the MBL in Woods Hole, MA, United States. Animals were maintained together in a 75-liter aquarium and a sand-filtered flow-thru seawater system. Tanks at OIST and at the MBL were open with no barriers to escape (Fig 1b, e). Animals typically began eating in lab within one to two days after collection and accepted freshly killed or store-bought frozen shrimp and crabs without training, in addition to live prey. Animals were surprisingly relaxed, and kept in tanks without lids or deterrents, as they generally did not attempt to escape.

### Identification

Animals were visually identified to species [28] and weighed. Sex identifications were also made based on male curling of the right third arm while moving, and on the presence of two large suckers at the proximal end of the arms in males but not females. For identification, octopus were tagged with silicone-based fluorescent elastomer *(Northwest Marine Technology)* that was injected into a small area in the dorsal mantle [2,31] (S2 Video). Anesthesia was not used, due to its potential negative influence on behavior in days after treatment, and due to the risk of losing animals. Injected octopus seemed lethargic immediately after injection but were back to normal in a few hours or the next morning. Experiments were not initiated until several days after injection to ensure all animals had recovered and were behaving normally.

### Experiments

Tagged animals were sorted into four groups of five to six animals, and cultured in four identical tanks, with mixed sizes and with sex balance, when possible. Tanks were circular, mostly opaque. Animals were maintained on an 11:13 hours light-dark cycle that roughly matched the local light cycle in Okinawa in late November and early December. The tanks were loosely covered with light-proof lids at the onset of the dark cycle to keep out most indoor lighting, but a very dim lighting was admitted by the plastic sides, roughly mimicking moon and stars light levels at night (Fig 1b). Similar to other nocturnal octopuses, *O. laqueus* are typically found in a den or sheltered space when not out foraging or mating during the day. Small clay pots (15 cm tall) were used as dens in the tanks, with each tank having either one pot per animal or one pot per three animals, depending on the treatment. Each pot had 4 large slits along the sides and a hole on top, allowing animals to freely and easily enter and leave the pots, and monitor activity outside the pot (Fig 1c). Sex-based social behaviors were also tested with each tank having either all females, all males, or an approximately equal mix. Clay pots were scored for animals three hours after the start of the light cycle (Fig 1d). Individuals were identified based on their elastomer tags when scoring clay pots for animals. After scoring, the animals were moved to small individual containers (Critter Keepers) with very small clay pots (5 cm tall) as dens, and the containers were returned to the larger tanks. One hour prior to the dark cycle, the small containers were moved to the bench top and two live or frozen shrimp or crab were added to each container. The animals were then returned to the main tank after a few hours. This ensured each animal was equally and adequately fed, and prevented fouling of the main tanks from left-over food, which rotted quickly in the warm conditions.

### Tests of social behavior

We examined the occupation patterns of pots in two sets of experiments, with varying number of animals in a tank and several mixing of sexes: 1) to test if *O. laqueus* prefer to be solitary versus social in a den, animals were placed in an open tank with one clay pot per octopus and five to six octopus per tank. Each tank included one to two large, three medium-sized, and one small animal. Two replicates of all females (Table 1a), all males (Table 1b), or an equal number of females and males (Table 2a,b) per tank were made. The number of octopus in each pot and in the tank outside the pots was assessed each morning for five to six days, with thirty-one tank assessments in total. 2) To test how social tolerance in *O. laqueus* is affected by limited den availability we used two tanks each having a total of six animals (and with 1:1 or 1:2 sex ratios in the tanks, due to limited number of animals) but only two clay pots per tank (Table 3a,b), with fourteen tank assessments in total. This allows one to see if animals preferred a densely populated clay pot over a less densely populated but exposed tank during the day.

**Table 1.**
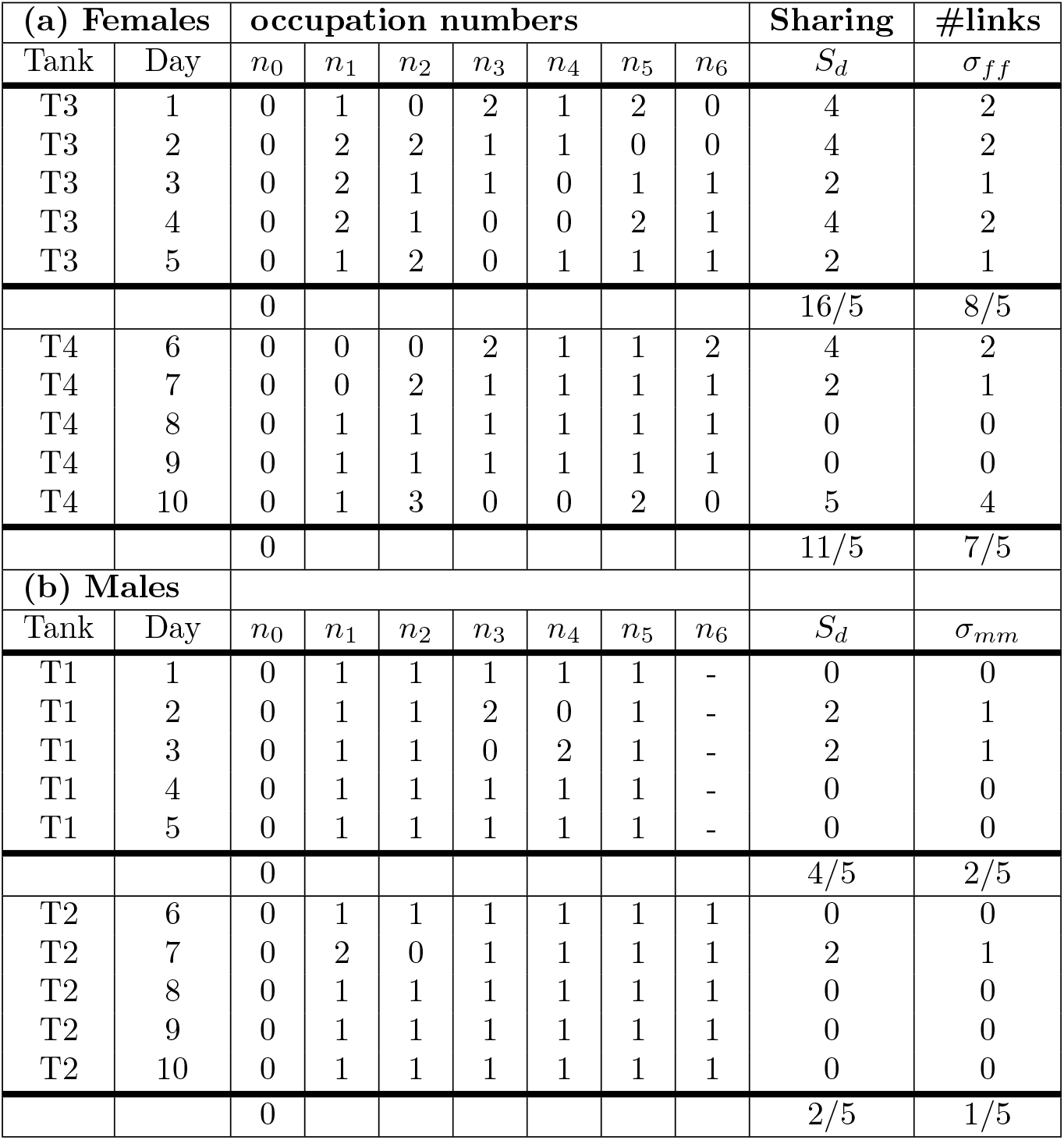
Occupation numbers for equal number of pots and animals (a) all females (b) all males. *n*_0_ indicates the number of outsiders, either females or males, that stay out of the pots.

**Table 2.** Occupation numbers for mixed sexes, *K* = 6 animals in *N* = 6 pots (*K_f_* = *K_m_* = 3).

| (a) |  | Female occ. numbers |  |  |  |  |  |  | Male occ. numbers |  |  |  |  |  |  | Sharing | #links |  |  |
| --- | --- | --- | --- | --- | --- | --- | --- | --- | --- | --- | --- | --- | --- | --- | --- | --- | --- | --- | --- |
| Tank | Day | $n_0$ | $n_1$ | $n_2$ | $n_3$ | $n_4$ | $n_5$ | $n_6$ | $n_0$ | $n_1$ | $n_2$ | $n_3$ | $n_4$ | $n_5$ | $n_6$ | $S_d$ | $\sigma_{ff}$ | $\sigma_{mm}$ | $\sigma_{fm}$ |
| T1 | 1 | 0 | 0 | 0 | 1 | 0 | 1 | 1 | 0 | 1 | 1 | 0 | 1 | 0 | 0 | 0 | 0 | 0 | 0 |
| T1 | 2 | 0 | 1 | 1 | 0 | 1 | 0 | 0 | 0 | 1 | 0 | 1 | 0 | 1 | 0 | 2 | 0 | 0 | 1 |
| T1 | 3 | 0 | 1 | 1 | 0 | 0 | 0 | 1 | 0 | 0 | 0 | 1 | 1 | 1 | 0 | 0 | 0 | 0 | 0 |
| T1 | 4 | 0 | 1 | 1 | 0 | 0 | 0 | 1 | 0 | 0 | 1 | 1 | 0 | 1 | 0 | 2 | 0 | 0 | 1 |
| T1 | 5 | 0 | 0 | 0 | 1 | 1 | 0 | 1 | 0 | 1 | 1 | 0 | 0 | 1 | 0 | 0 | 0 | 0 | 0 |
| T1 | 6 | 0 | 1 | 0 | 0 | 0 | 0 | 2 | 0 | 0 | 1 | 1 | 1 | 0 | 0 | 2 | 1 | 0 | 0 |
|  |  | 0 |  |  |  |  |  |  | 0 |  |  |  |  |  |  | 6/6 | 1/6 | 0 | 2/6 |
| (b) |  |  |  |  |  |  |  |  |  |  |  |  |  |  |  |  |  |  |  |
| T3 | 1 | 1 | 1 | 0 | 1 | 0 | 0 | 0 | 0 | 1 | 0 | 0 | 0 | 1 | 1 | 2 | 0 | 0 | 1 |
| T3 | 2 | 0 | 0 | 1 | 0 | 1 | 1 | 0 | 0 | 0 | 0 | 1 | 0 | 1 | 1 | 2 | 0 | 0 | 1 |
| T3 | 3 | 0 | 0 | 0 | 0 | 1 | 1 | 1 | 0 | 0 | 1 | 1 | 0 | 1 | 0 | 2 | 0 | 0 | 1 |
| T3 | 4 | 0 | 0 | 1 | 0 | 0 | 0 | 2 | 0 | 1 | 1 | 0 | 0 | 1 | 0 | 4 | 1 | 0 | 1 |
| T3 | 5 | 0 | 0 | 1 | 1 | 0 | 0 | 1 | 0 | 1 | 0 | 1 | 0 | 0 | 1 | 4 | 0 | 0 | 2 |
|  |  | 1/5 |  |  |  |  |  |  | 0 |  |  |  |  |  |  | 14/5 | 1/5 | 0 | 6/5 |

**Table 3.** Occupation numbers for mixed sexes, *K* = 6 animals in *N* = 2 pots. (a) *K_f_* = *K_m_* = 3 (b) *K_f_* = 4, *K_m_* = 2.

| (a) |  | Females |  |  | Males |  |  | Sharing | #links |  |  |
| --- | --- | --- | --- | --- | --- | --- | --- | --- | --- | --- | --- |
| Tank | Day | $n_0$ | $n_1$ | $n_2$ | $n_0$ | $n_1$ | $n_2$ | $S_d$ | $\sigma_{ff}$ | $\sigma_{mm}$ | $\sigma_{fm}$ |
| T2 | 1 | 1 | 1 | 1 | 0 | 1 | 2 | 5 | 0 | 1 | 3 |
| T2 | 2 | 0 | 1 | 2 | 0 | 1 | 2 | 6 | 1 | 1 | 5 |
| T2 | 3 | 1 | 1 | 1 | 0 | 1 | 2 | 5 | 0 | 1 | 3 |
| T2 | 4 | 1 | 1 | 1 | 0 | 2 | 1 | 5 | 0 | 1 | 3 |
| T2 | 5 | 1 | 0 | 2 | 0 | 2 | 1 | 5 | 1 | 1 | 2 |
| T2 | 6 | 1 | 1 | 1 | 1 | 1 | 1 | 4 | 0 | 0 | 2 |
| T2 | 7 | 0 | 1 | 2 | 0 | 1 | 2 | 6 | 1 | 1 | 5 |
|  |  | 5/7 |  |  | 1/7 |  |  | 36/7 | 3/7 | 6/7 | 23/7 |
| (b) |  |  |  |  |  |  |  |  |  |  |  |
| T4 | 1 | 1 | 1 | 2 | 0 | 0 | 2 | 4 | 1 | 1 | 4 |
| T4 | 2 | 1 | 1 | 2 | 0 | 1 | 1 | 5 | 1 | 0 | 3 |
| T4 | 3 | 1 | 2 | 1 | 0 | 1 | 1 | 5 | 1 | 0 | 3 |
| T4 | 4 | 1 | 1 | 2 | 0 | 1 | 1 | 5 | 1 | 0 | 3 |
| T4 | 5 | 1 | 1 | 2 | 0 | 1 | 1 | 5 | 1 | 0 | 3 |
| T4 | 6 | 0 | 2 | 2 | 1 | 1 | 0 | 5 | 2 | 0 | 2 |
| T4 | 7 | 0 | 2 | 2 | 1 | 0 | 1 | 5 | 2 | 0 | 2 |
|  |  | 5/7 |  |  | 2/7 |  |  | 34/7 | 9/7 | 1/7 | 20/7 |

Methods to perform the social behavior experiments are detailed here: http://dx.doi.org/10.17504/protocols.io.w9nfh5e

## Results

*O. laqueus* was surprisingly abundant at Maeda Flats in Okinawa during the fall, winter, and early spring. Animals were common in holes or dens in the sand and reef rubble (Fig 1a) that were within a few meters of one another, suggesting that a given individual is likely to encounter multiple other individuals on a given night of foraging and hunting. On three occasions while diving or intertidal walking, sets of two octopus were in dens or holes that were close enough for the animals to touch one another, and it was possible that they were sharing a single den with multiple entrances (S1 Video). The abundance and proximity of *O. laqueus* in the field raised the possibility that it is a social octopus, leading us to ask the specific question, is *O. laqueus* tolerant of other octopus under dense conditions in a shared tank or even den, and to design a series of experiments to test this question.

To initially test if *O. laqueus* might be socially tolerant, ten or more octopus were placed in ten-liter buckets over the course of collection in the field to see how they reacted. Octopus were left in buckets for one to four hours and checked periodically. They did not exhibit any obvious aggressive interactions beyond occasional color flashes due to disturbances from researchers with flashlights and would often sit in contact with one another. In addition, and in contrast to other species in the same habitat, like *Abdopus aculeatus* and *Octopus incella*, animals rarely attempted to escape out of the open buckets, despite the stress of collection, dense conditions, and limited seawater.

*O. laqueus* were easy to care for in group cultures in lab at densities up to one animal per 15 liters, and up to 15 animals in a 250-liter tank, with a mix of sexs, and with a mix of small to large sizes for the species. The animals were surprisingly relaxed, and were kept in tanks without lids or deterrents, as they generally did not attempt to escape. Out of over 100 animals brought into the lab over several years, only a few ever escaped or disappeared from their tank. At least one incident of escape appeared to be related to poor water conditions that unexpectedly arose, while another might have been due to its extremely small size, as it was much smaller than any other animals brought in.

To test how long animals might be group cultured in open tanks in a lab environment, three small juvenile animals (one male and two females, each around 10 grams) were shipped from Okinawa, Japan and maintained at the Marine Biological Station in the United States in a single 50-gallon tank that was open without a lid or other deterrents. The animals did well in group culture for over four months and did not try to escape. One animal laid eggs after 3 months but disappeared a few days after laying, possible due to cannibalism related to her brooding embryos. A second female laid eggs 4 months after arrival. To ensure the remaining male did not endangered the mom or her embryos, the male was moved to a separate tank the following day, and was later euthanized for another experiment. The mom successfully cared for her embryos through hatching (Fig 1f) and died of natural causes after five months.

*O. laqueus* were kept in aquaria with clay pots serving as dens. At night, octopus were observed roaming their tanks, hunting and eating prey, and interacting. Each morning, octopus would select a pot to stay in during the day, or rarely remain outside the pots in the tank. Multiple individuals were often found co-occupying a single pot, within arm’s reach or in contact with one another inside the pot, suggesting *O. laqueus* will share a den in the lab.

The daily pot occupancies, observed over 45 days, are shown in Tables 1–3. Each table specifies the number of available pots *N*, the number of females *K_f_* or males *K_m_* in the tank (*K_m_* + *K_f_* = *K*), the number of animals that were found each day inside pot *i, n_i_* (*i* = 1,…, *N*), and the number animals that stayed in the open space outside the clay pots, *n*_0_, so that in total 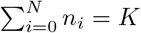.

Pot occupancy across 31 experiments having an equal number of octopus and pots (*N* = *K*) ranged from zero to three animals in a pot. Five occupancy patterns were observed per tank per day: 1) each pot in tank contained a single octopus, 2) one pot contained two octopus and remaining pots in a tank contained one or zero octopus, 3) one pot contained two octopus, remaining pots in a tank contained one or zero octopus, and the tank itself contained one octopus, 4) two pots contained two octopus and remaining pots in a tank contained one or zero octopus, 5) one pot contained three octopus, one pot contained two octopus, and remaining pots in a tank contained one or zero octopus (Tables 1–2). The total number of sharing animals per day, 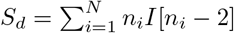 (*I* indicates summing over *n_i_* ≥ 2), ranged from zero to five. Specifically, we found that *S_d_* ≥ 2 i.e., at least two animals were sharing a pot, in 19 out of 31 days (61%). Then, looking at the subset of *N* = *K* = 6 (omitting the all-male *N* = 5 replicas in Table 1b, and a single incident with *n*_0_ ≠ 0 in Table 2b), we found that *S_d_* ≥ 2 in 16 out of 25 days (64%). These numbers are sufficiently high to demonstrate that *O. laqueus* are not totally solitary and can be tolerant of sharing a clay pot or den with one or more individuals. At the same time, it’s clear that the animals are far from being neutral (indifferent to the presence of others) because, for *K* independent animals distributed among *N* jars, the probability of non-sharing would be *P*(*S_d_* = 0) = *K*!*N*^-*k*^ = 1.5%. (Note that if, in addition, the animals can be viewed as identical and indistinguishable, this probability is reduced to *P*(*S_d_* = 0) = *K*!(*N* – 1)! /(*N* + *K* – 1)! = 0.22%). Averaging the occupation numbers of all the pots over 25 days, we also found that 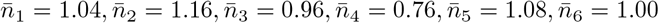, verifying thereby that all pots are statistically identical (the deviations compared to the expected average value of 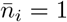 are insignificant). Thus, clay pot or den selection is not an entirely random process, and there is an anti-social behavioural component at play, keeping pot sharing at levels lower than predicted by a neutral random model.

Sex analysis of the pot occupancy data suggests that the anti-social behavior component is coming primarily from male-male interactions but occurs at statistically significant levels even in all-female tanks. Indeed, the average sharing number of all-females configurations is 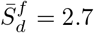 whereas the all-males average sharing is 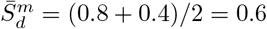. Therefore, females are much friendlier than males, however, both sexes are less friendly than neutral animals (for comparison, the average sharing level, assuming *K* = 6 independent animals distributed in *N* = 6 jars, is 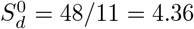. Furthermore, considering the case of mixed sexes (Table 2), one can verify that most of the sharing events occurred by forming female-male pairs. This (rather unsurprising) tendency persists even in the case of limited number of dens, *N* < *K* (Table 3).

The pot occupancies across all the experiments having more octopus than pots (*N* < *K*) ranged from two to four animals in a pot (Table 3), and the daily sharing numbers ranged accordingly from four to six sharing animals per day. We found that, out of fourteen tank examinations, *S_d_* = 4, 6 each occurred twice and *S_d_* = 5 occurred 10 times (71%). However, limiting dens also increased the number of ‘outsiders’ (namely those animals, either females or males, that stay in the open environment outside the pots) so that a solitary octopus was found outside the pots on most days. Specifically, we found that *n*_0_ = 0 occurred only twice (14.3%), *n*_0_ = 1 occurred 11 times (78.6%), and *n*_0_ = 2 happened once (7.1%). Thus, overall, limiting dens increased the amount of social sharing but, at the same time, forced some fraction of the animals to stay out of the dens.

To further quantify the pot occupancies, we calculated the observed number of pairwise links formed daily by the animals:

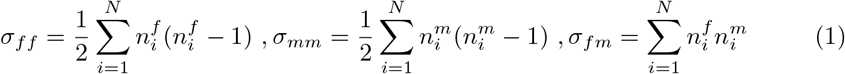

The results, measured over the total of 45 days are presented in Tables 1–3. The mean number of links, averaged over *M* days for various densities of animals and mixtures of sexes, are summarized in Table 4.

**Table 4.**
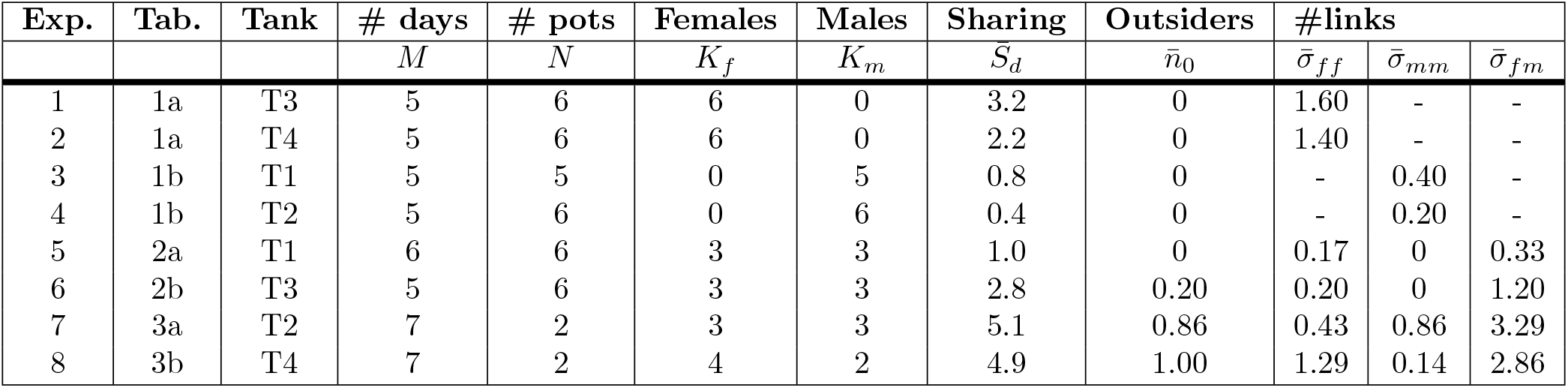
Summary of measurements for eight experimental setups showing the sharing number 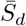, number of outsiders 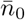, and the number of links 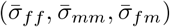, averaged over observations.

## Modelling

In the following we obtain a statistical description of pot occupancies. We employ the standard statistical mechanics approach, in which distributions and correlations – such as the probability distribution of sharing, *P*(*S_d_*) are derived from a Hamiltonian under the maximum entropy principle [32]. Maximum entropy then leads to the least structured model that is still consistent with the empirical observations. Our model might be called the ‘‘housemate model:” you will let me share your home only if I get along with everyone else in the house; similarly, everyone already living there must get along with one another. Such pairwise propensities, affinities, or proclivities are taken to be independent of and uncorrelated with one another. The probability that *N* agents get along with each other decays exponentially with the total number of pairwise interactions between, or distinct pairs of, agents, *N*(*N* – 1)/2. Of course the model is simplistic, but with readily achievable values of *N*, it could be misleading to try to fit a more complex model.

For simplicity, let us start with the case of a single sex distribution. Assuming *K* identical (indistinguishable) animals distributed among *N* pots, any daily configuration of animals is completely determined by specifying a set of occupation numbers:

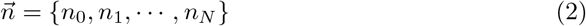

The mean number of outsiders averaged over *M* days, 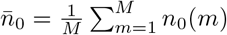, and the mean number of pair-interactions, 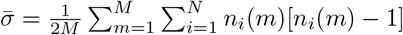, are measured experimentally (see Table 4). The probability distribution, 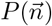, that maximises the entropy 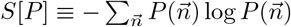 under the empirical constraints 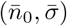 is the canonical distribution 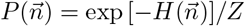, where

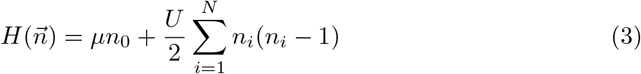

is the Hamiltonian and *Z* is the partition function, obtained by summing over all configurations, 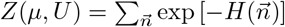. Eq (3) resembles the bosonic Hubbard model which is well known in condensed matter physics [33,34]. The parameters introduced in (3) are the on-site interaction *U* (which can be attractive, *U* < 0, or repulsive, *U* > 0), and the chemical potential *μ* which penalizes the outsiders (*μ* > 0). These parameters are found by imposing the conditions

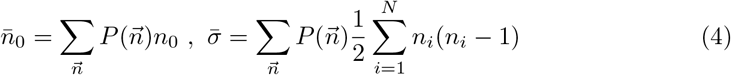

so that empirical-averaging coincides with ensemble-averaging with respect to 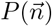. Namely,

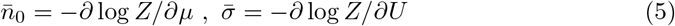

Alternatively, the parameters can be obtained by the maximum likelihood condition,

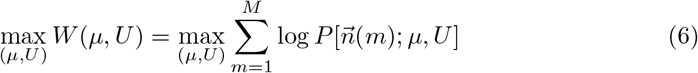

where 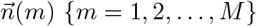 are the observed configurations, and log *P* = – (*H* + log *Z*). The error in estimating the parameters is then given by the Gaussian fluctuation [35] (a.k.a. Fisher information matrix), evaluated at the maximum-likelihood solution (*μ*_0_, *U*_0_):

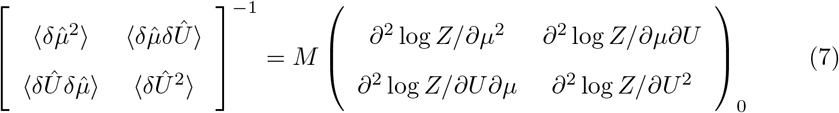

For *n*_0_ ≡, *μ* is traced out of Eq (3) (i.e., *μ* → ∞) so that the partition function is independent of the chemical potential, *Z* = *Z*(*U*). As a result, Eq (7) reduces to

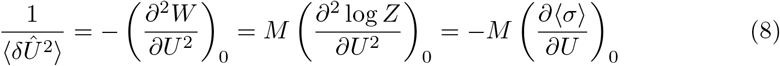

The linear-response term on the RHS of (8) is related to the variance of σ by the fluctuation-dissipation theorem [36]. Thus, 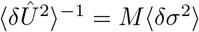.

### Modeling equal number of octopus and pots

The total number of configurations for *K* identical animals occupying *N* pots is

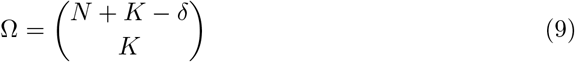

where *δ* ≡ if 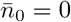 and zero otherwise (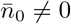 means that the animals can dwell somewhere in the tank outside the pots. Combinatorically, this amounts to having an additional available ‘slot’). Referring to the first four rows in Table 4, with *N* = *K* = 6 and *δ* =1, the number of configurations is Ω = (2*N* – 1)!/[*N*!(*N* – 1)!] = 462. The calculation of the partition function is, therefore, amendable to numerical computation. Note that distinguishable animals would have assumed a total of *N^k^* = 46656 configurations which is also doable.

The partition function (more precisely, the free energy *F* – log *Z*) as a function of *U* is plotted in Fig 2. Therefore, given *F*(*U*) and solving 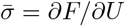 [Eq (5)] for *U*, we find that

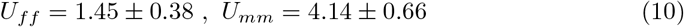

**Fig 2.**
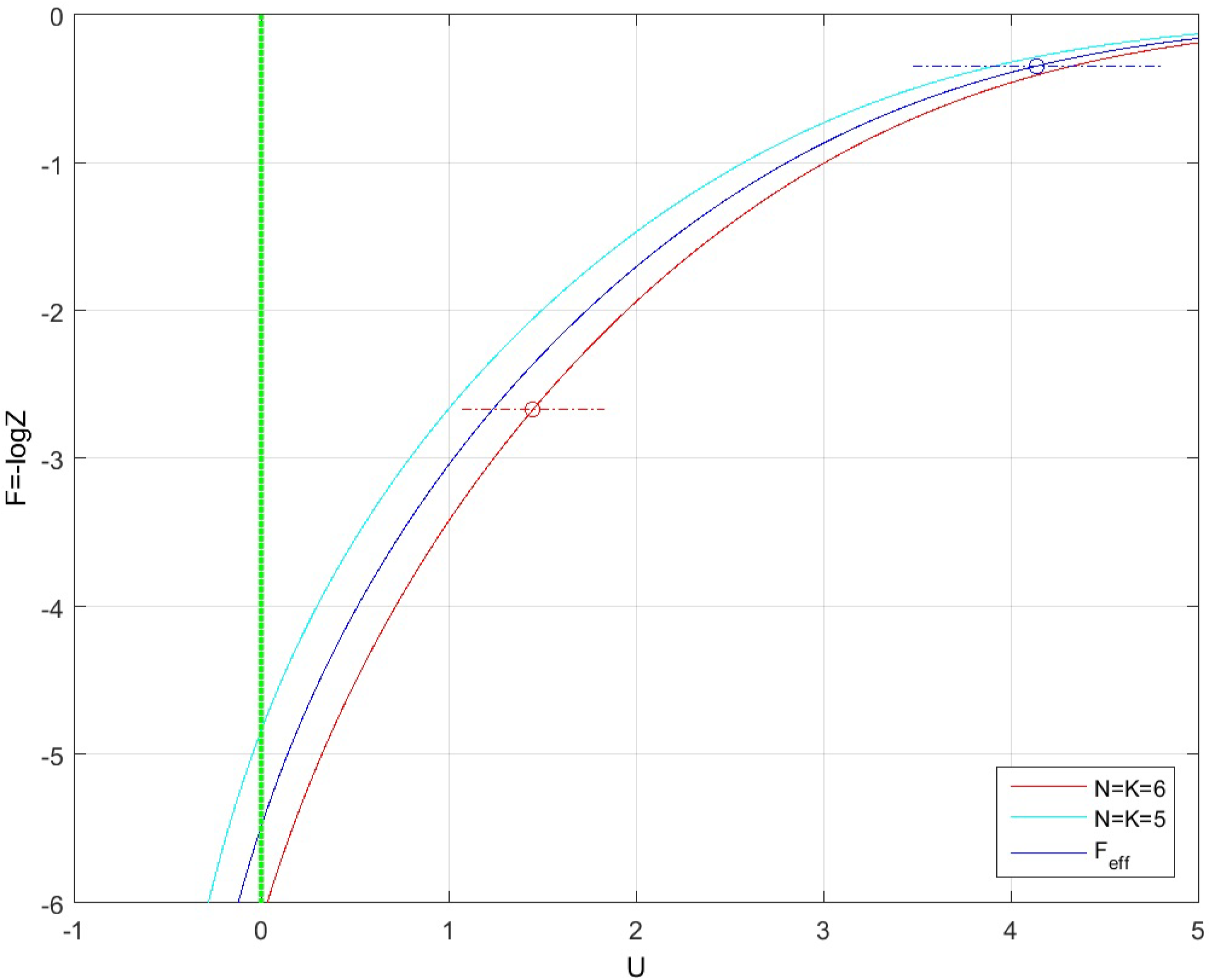
The free energy, *F* = – log *Z*, as a function of the interaction parameter *U*. (a) for *N* = *K* = 6 (b) for *N* = *K* = 5. (c) The combined free-energy 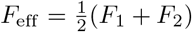. The estimated values of the interaction, *U_ff_* and *U_mm_* [Eq. (10)], are shown together with their corresponding error-bars. Both females and males are far from being neutral (green line, *U* = 0). In a single-sex environment females are more social than males.

As expected, in a non-mixed environment females are friendlier than males. However, compared to neutral animals (*U* = 0) both sexes exhibit significant repulsive interaction. The *t*-statistic for the difference between sexes is 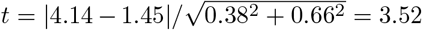 (with a *p*-value= 0.002). These results are based on combining two replicates consisting of a total of 10 measurements for each sex (see Table 1). For males, since *N*_1_ = *K*_1_ = 6 and *N*_2_ = *K*_2_ = 5, the combined free energy for two replicates is given the weighted average, *F*_eff_ = (*M*_1_*F*_1_ + *M*_2_*F*_2_)/(*M*_1_ + *M*_2_). In this case, Eq (5) takes the form: 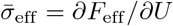, where 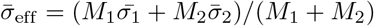 is the effective number of links. We computed the following quantities as a function of the interaction parameter *U* (Fig 3): (i) the average number of links 〈σ〉 = *∂F*/*∂U* (ii) the canonical distribution 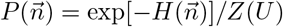 and the log-likelihood function 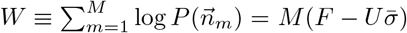 and (iii) the fluctuation 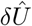 according to Eq (8).

**Fig 3.**
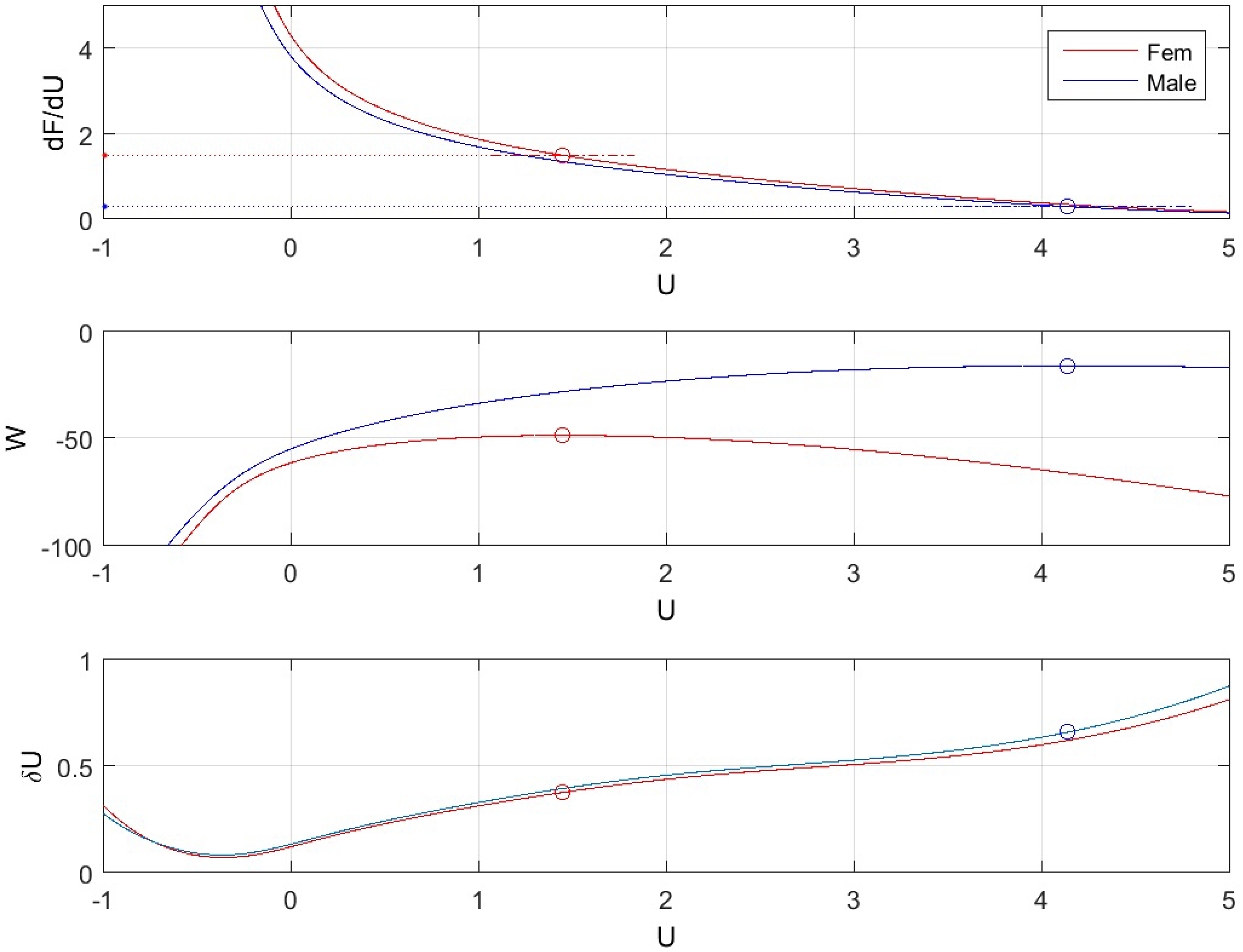
(a) The average number of links 〈σ〉 = *∂F/∂U* and (b) the log-likelihood function. *W* = *F* – *U*〈σ〉 showing that *W* assumes its maximal value when 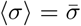. (c) the fluctuation 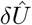. (solid-red line: females, solid-blue line: males).

As a consistency check of the model, we’ve also calculated the average sharing-number in terms of the canonical distribution 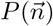. Namely, 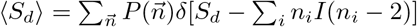. We found that the average sharing numbers, calculated at the corresponding maximum likelihood solutions (10) (i.e., *U_ff_* – 1.45, *U_mm_* = 4.14), are

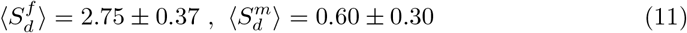

These values are in good agreement with the experimental results, 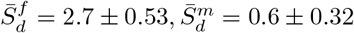. In particular, the estimated errors in (11) are smaller than the empirical ones and, as shown in Fig 4, the empirical values lay well within the estimated confidence levels. Since, 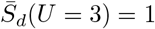 (Fig 4), it follows that, for *U* ≤ 3 one typically observes at least one pot with sharing animals, whereas for *U* > 3 sharing is much suppressed. Also note that both values in (11) differ significantly from the expected sharing level of neutral animals:

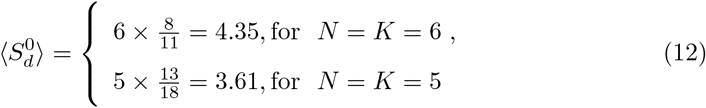

**Fig 4.**
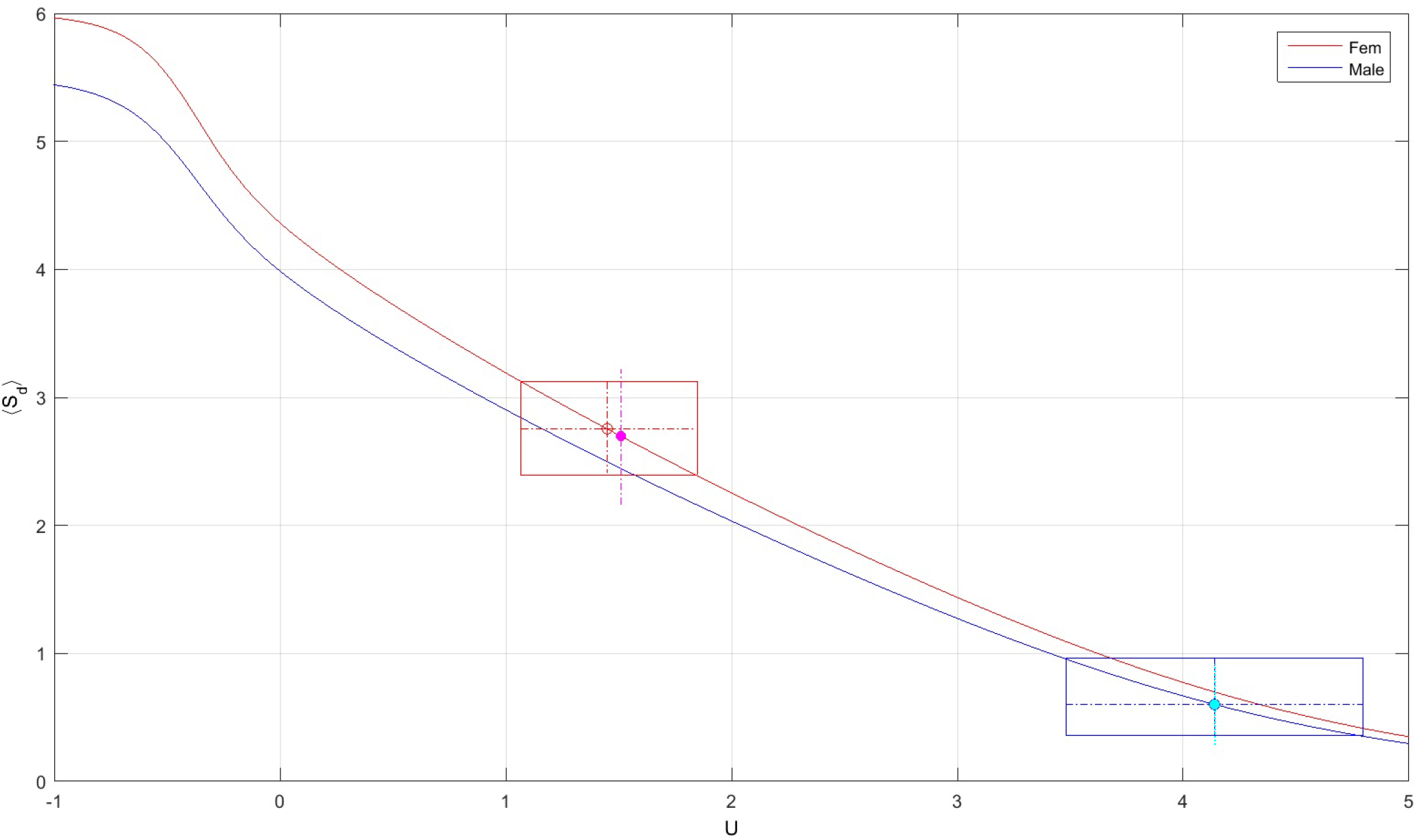
The average sharing number. 〈*S_d_*〉 as a function of *U* for females (red) and males (blue). Females: 〈*S_d_*〉 evaluated at *U_ff_* = 1.45 gives 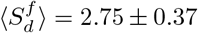 (indicated by the red-rectangle) and the experimental value is 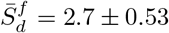 (dotted-magenta). Males: 〈*S_d_*〉 evaluated at *U_mm_* = 4.14 gives 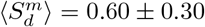 (blue-rectangle) and the experimental value is 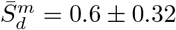 (dotted-cyan).

The full sharing distribution as a function of the interaction parameter, *P_S_d__* (*k|U*) {*k* = 0, 2,…, *K*}, is shown in Fig 5 (for *N* = *K* = 6). We find that the non-sharing probability *P*_0_ ≡ *P_S_d__*(*k* = 0|*U*), evaluated at the saddle points (10), is 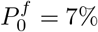 for females and 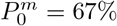 for males. Clearly, both values are larger than the non-sharing probability of neutral animals (either indistinguishable ones for which 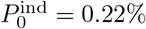, or distinguishable ones with 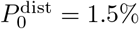). More generally, we examined the Kullback-Leibler distance between the empirical sharing distribution, 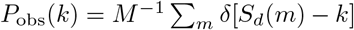, and the probability *P_S_d__* (*k|U*) calculated as a function of *U* by using the distribution function 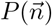. We found (Fig 6), that the KL-distance *D*[*P*_obs_(*S_d_*)||*P*_cal_(*S_d_*|*U*)] assumes its minimal value – respectively for females and males, at *U* = (1.44, 4.34) which is again very close to the saddle points (10). Remarkably, this holds even though the number of observations, *M* = 10, is pretty small. In addition, the one-parameter model 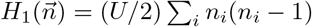 (resulting from Eq 3) by setting *μ* → ∞) has the smaller AIC as compared other polynomial models (see Table 5 and Fig 7).

**Table 5.** Comparison of AIC (Akaike Information Criterion) for 3 polynomial models with K=6 females in N=6 pots, measured over M=10 days. *AIC* = –2log *L* + 2*p* + 2*p*(*p* + 1)/(*M* – *p* – 1), BIC=—2log *L* + 2*p* log *M*, where *p* = (1, 2, 3) is the number of parameters.

| Model |  | parameters | loglike | AIC | BIC |
| --- | --- | --- | --- | --- | --- |
| Hubbard | $H_1(\vec{n}) = (U/2) \sum_i n_i(n_i - 1)$ | $U = 1.44$ | -48.4135 | 99.3269 | 101.4321 |
| Linear | $H_2(\vec{n}) = (V/2) \sum_i n_i I[n_i - \nu]$ | $V = 0.98, \nu = 2$ | -50.5966 | 106.9075 | 110.4036 |
| 3rd order polynomial | $H_3(\vec{n}) = (1/2) \sum_i W_i(n_i)$ | $W(0) = 0, W(1) = 0.04,$<br>$W(2) = 1.63, W(n \geq 3) = 4.32$ | -48.3707 | 106.7413 | 110.5568 |

**Fig 5.**
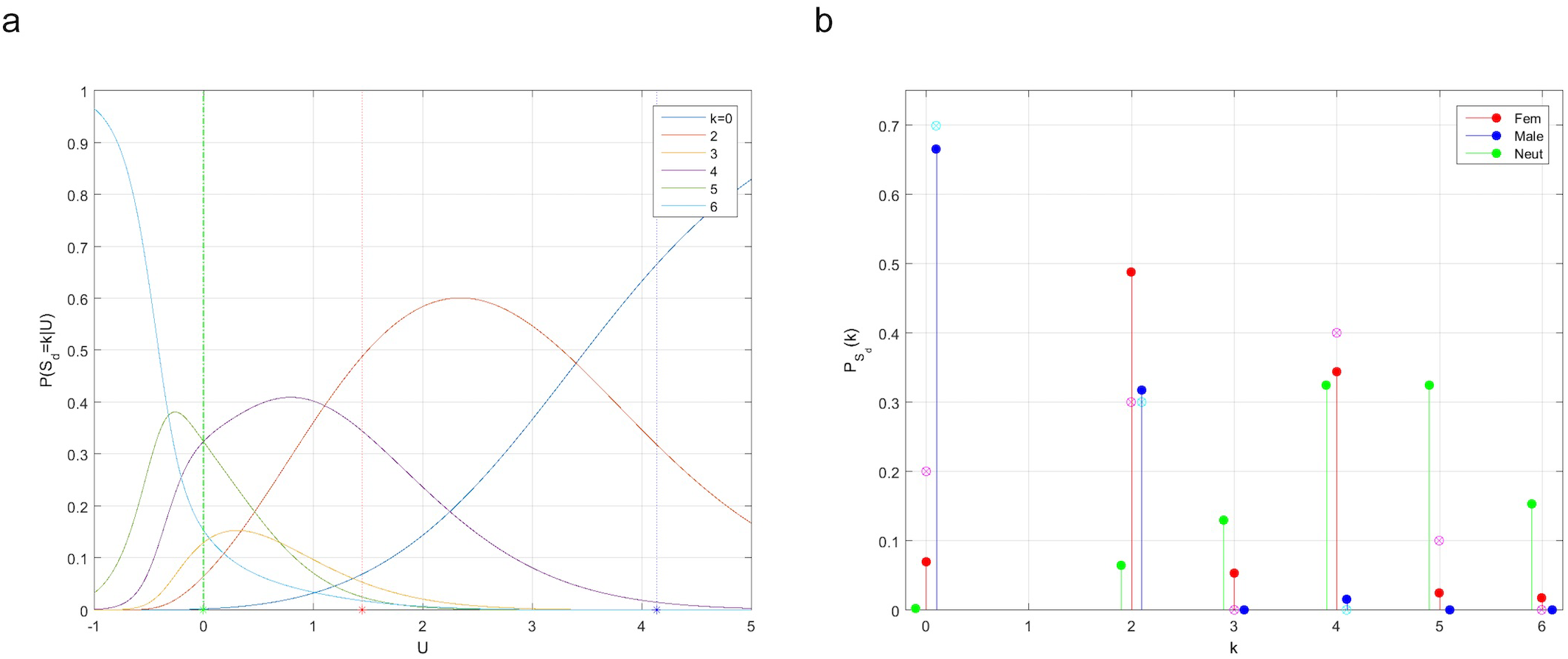
The sharing distribution as a function of the interaction parameter, *P_S_d__* (*k|U*) {*k* = 0, 2,…, *K*} for *K* = 6 animals in *N* = 6 jars. (a) The probability of non-sharing (blue solid line) is 7% for females (red dot at *U* = 1.45) and for males 67% (blue dot at *U* = 4.14). For neutral animals the probability of non-sharing is 1.5% (green dot at *U* = 0). (b) Comparison of the sharing probability for females (red), males (blue) and neutral animals (green). The empirical probability, obtained by averaging of 10 days, is also shown for females (magenta) and males (cyan).

**Fig 6.**
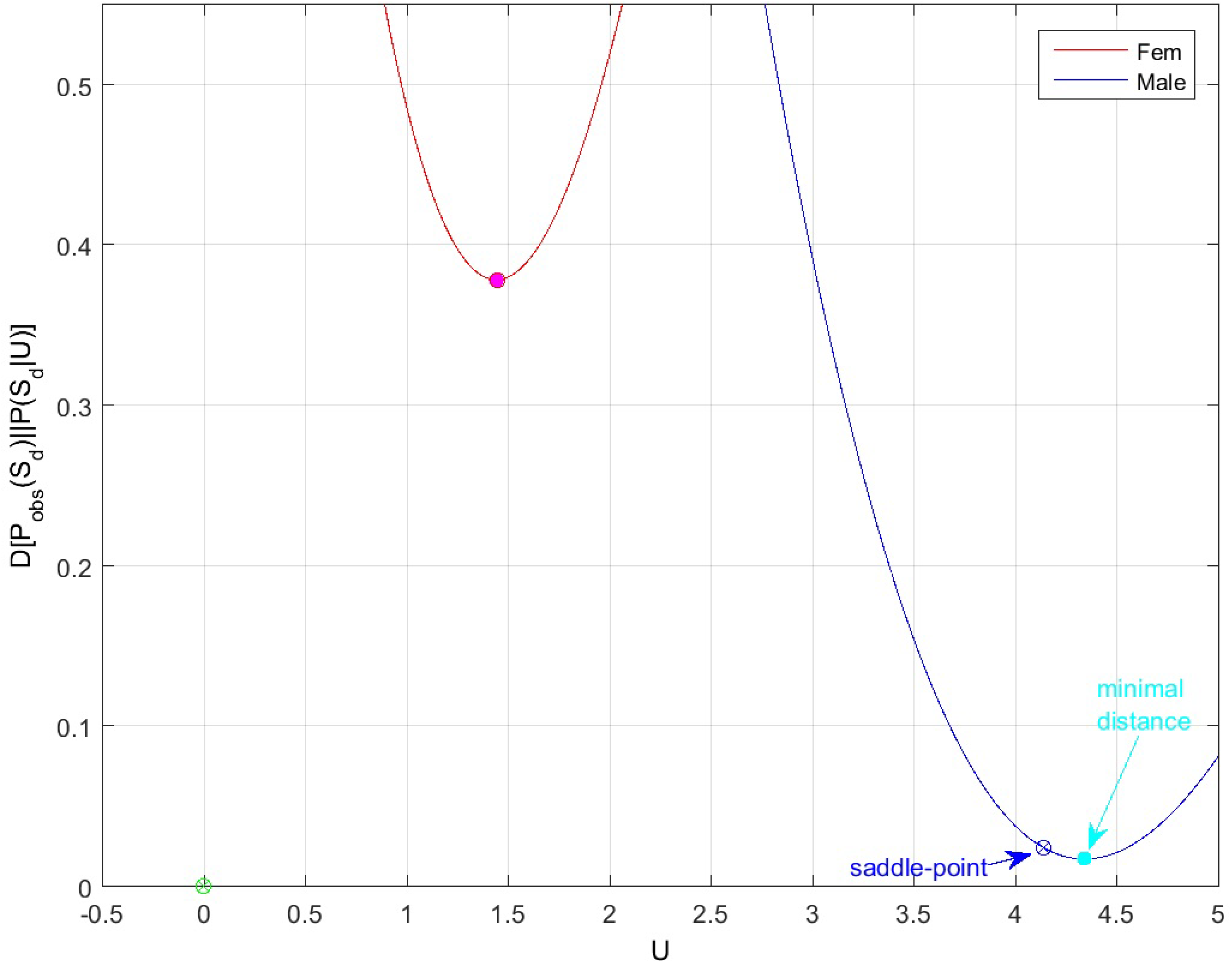
(a) **The KL-distance** *D*[*P*_obs_(*S*_d_)||*P*(*S*_d_|*U*)], between the empirical sharing distribution and the calculated sharing distribution as a function of U, for females (red) and males (blue).

**Fig 7.**
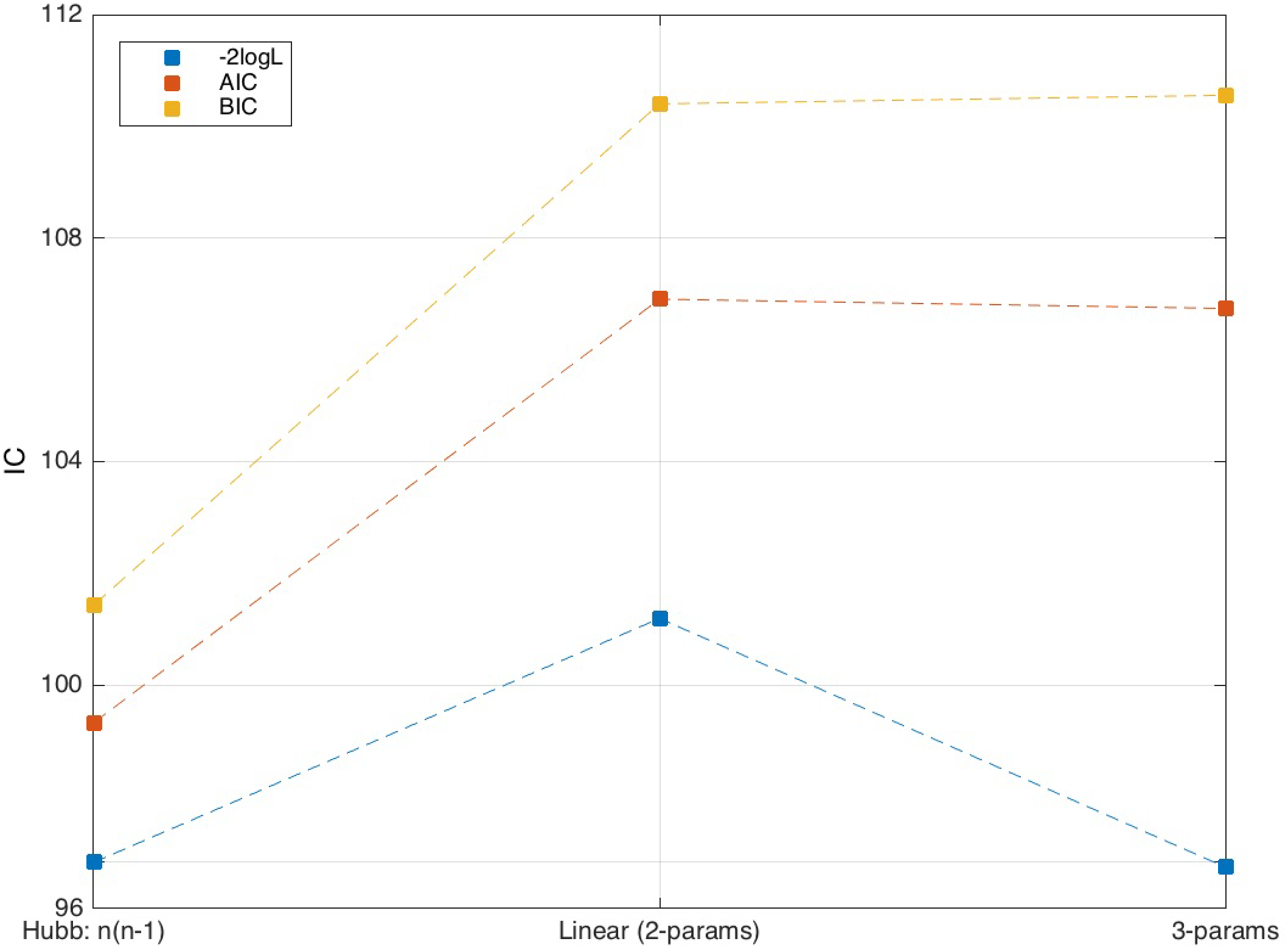
Comparison of AIC (Akaike Information Criterion) for 3 polynomial models with *K* = 6 females in *N* = 6 pots. (a) Hubbard: 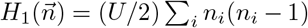. (b) linear 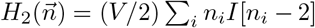. (c) 3-parameters: 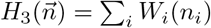, with *W_i_*(0) = 0, *W_i_*(1) = *ω*_1_, *W_i_*(2) = *ω*_2_, *W_i_*(*n_i_* ≥ 3) = *ω*_3_. The maximum likelihood L is obtained, respectively, for *U* = 1.44, *V* = 0.98 and *ω* = (0.04,1.63,4.32). Here AIC=-2log*L* + 2*p* + 2*p*(*p* + 1)/(*M* -*p* – 1), BIC=-2 log *L* + 2*p*log *M*, where *p* = (1, 2, 3) is the number of parameters and *M* = 10 is the numbers of measurements.

### Modeling more octopus than pots

The model of Eq (3) can be easily extended for describing experiments with mixed sex/species (as long as distinct species can share a spot without eating one another):

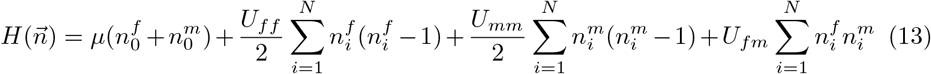

Here 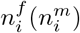 is the number of females(males) occupying pot *i* (out of *N*) and 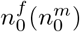 is the number of outsiders. This model allows one to consider different interactions between sexs. For example, *U_fm_* ≤ 0 ≤ *U_ff_* ≤ *U_mm_* would describe a kind of ‘straight’ animals, having attractive inter-sex interaction and repulsive interaction within the same sex, with the females being more social than the males. The total number of configurations associated with mixed populations as in (13) is

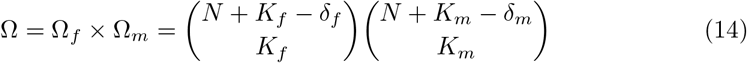

where 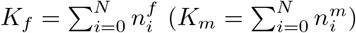 is number of females (males) and 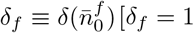, if 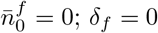 otherwise; and similarly for 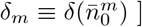.

We first applied the extended model (13) to the case of mixed sexs with equal number of animals and pots: *K_f_* = *K_m_* = 3,*N* = *K_f_* + *K_m_* = 6. Referring to Table 4, we find that *δ_f_* =0 and 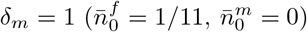. The number of configurations is then 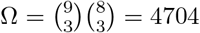. Combining tanks #1 and #3, we also observed that 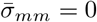 (males never shared pots with any other males for 11 days). Therefore, tracing out *U_mm_* and solving *dF/∂μ* = 1/11, *dF/dU_ff_* = 2/11, *∂F/∂U_fm_* = 8/11, we find that

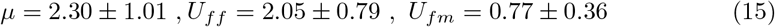

The female-female interaction is consistent with the previous result [Eq (10)] obtained in a single-sex environment. The chemical potential *μ* being of the same order of magnitude as *U_ff_* is sufficient to prevent females from staying outside the pots. The female-male interaction *U_fm_* is much less repulsive than either Uff or *U_mm_*. The error estimates in (15) are obtained, as in (7), by calculating the Gaussian fluctuation of the free-energy at the maximum-likelihood solution. Thus, introducing *x* (*μ, U_ff_,U_fm_*),

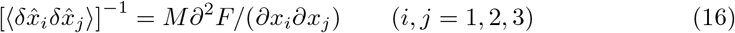

Next, we considered the case of dense pots *N* = 2 < *K_f_* + *K_m_* = 6 For tank #4, containing 4 females and 2 males that are sharing 2 pots, Eq (14) gives 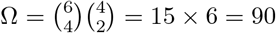 configurations. For tank #2, with 3 females and 3 males 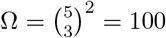. Such small number of configurations enables one to obtain the exact partition function and infer the four coupling constants of 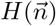. In practice however, the number of samples *M* = 7 is also very small, so the expected accuracy of these parameters is rather low. The results are summarised in Table 6. Tank #4 looks promising: females are more social than males and *f* – *m* interaction is on the verge of attraction

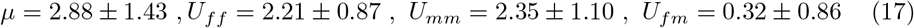

On the other hand, in tank #2 the males look more social than females:

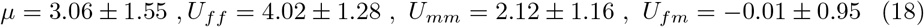

**Table 6.** The estimated interaction parameters for eight experimental setups and their combinations.

| Exp. | Tab. | Tank | # days | # pots | Fem | Male | # config | interaction |  |  |  |
| --- | --- | --- | --- | --- | --- | --- | --- | --- | --- | --- | --- |
| | | | $M$ | $N$ | $K_f$ | $K_m$ | $\Omega$ | $\mu$ | $U_{ff}$ | $U_{mm}$ | $U_{fm}$ |
| 1 | 1a | T3 | 5 | 6 | 6 | 0 | 462 | - | $1.31 \pm 0.51$ | - | - |
| 2 | 1a | T4 | 5 | 6 | 6 | 0 | 462 | - | $1.60 \pm 0.56$ | - | - |
| 1+2 | | | 10 | 6 | 6 | 0 | 462 | - | $1.45 \pm 0.38$ | - | - |
| 3 | 1b | T1 | 5 | 5 | 0 | 5 | 126 | - | - | $3.52 \pm 0.85$ | - |
| 4 | 1b | T2 | 5 | 6 | 0 | 6 | 462 | - | - | $4.84 \pm 1.09$ | - |
| 3+4 | | | 10 | 5,6 | 0 | 5,6 | 462 | - | - | $4.14 \pm 0.66$ | - |
| 5 | 2a | T1 | 6 | 6 | 3 | 3 | 3136 | - | $2.47 \pm 1.10$ | - | $1.60 \pm 0.70$ |
| 6 | 2b | T3 | 5 | 6 | 3 | 3 | 4704 | $1.25 \pm 1.03$ | $1.66 \pm 1.11$ | - | $0.18 \pm 0.43$ |
| (a) 5+6 | | | 11 | 6 | 3 | 3 | 4704 | $2.30 \pm 1.01$ | $2.05 \pm 0.79$ | - | $0.77 \pm 0.36$ |
| (b) 7 | 3a | T2 | 7 | 2 | 3 | 3 | 100 | $3.06 \pm 1.55$ | $4.02 \pm 1.28$ | $2.12 \pm 1.16$ | $-0.01 \pm 0.95$ |
| (c) 8 | 3b | T4 | 7 | 2 | 4 | 2 | 90 | $2.88 \pm 1.43$ | $2.21 \pm 0.87$ | $2.35 \pm 1.10$ | $0.32 \pm 0.86$ |

This ‘anomaly’ can be traced back to a high degree of individual variety (it turns out that a certain large female, named *2RG*, sits most of the time out of the pots and seems to be extremely anti-social).

All the experimental results can be treated on the same footing by combining the interaction parameters, obtained separately under different experimental conditions, into a single set of properly weighed parameters. Referring to Table 6 and combining together the results of five different setups, (1+2), (3+4), (5+6), (7) and (8) [see also Eqs. (10),(15),(17),(18)], we find:

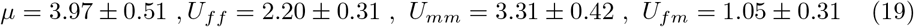

Eq (19) specifies the most probable set of interaction parameters that are consistent with the total of 45 available measurements. These values can be used in 2 ways: first, for identifying potential outliners and second, for the prediction of the behavior over a large set of experimental designs. As an example, let’s consider *K* = 6 animals distributed among a varying number of pots *N* = (1, 2,…,8) with several possible mixtures of sexs, *K_f_* =0,1,…, 6 (*K_m_* = *K* – *K_f_*). In this case, all quantities of interest, such as the number of outsiders *n*_0_ or the female-male linkage *σ_fm_* (which may well affect factors like potential mating, rate of cannibalism etc.), are determined by two parameters: the specific volume *N/K* and the sex mixture *K_f_*/*K*.

In Figs 8a-b, 〈*n*_0_〉, 〈σ_fm_〉 are shown as functions of *N* and *K_f_*. As expected, both 〈*n*_0_〉 and 〈*σ_fm_*〉 assume their maximal values when the number of pots is limited (*N* = 2) and the mixture of sexs is balanced (*K_f_* = *K_m_*). Fig 8 suggests that the two empirical points (b, c), described by Eqs. (17) and (18), lay reasonably close to the respectively calculated curves. On the other hand, the point (a) corresponding to Eq (15), forms a clear cut ‘outliner’. This discrepancy can be attributed to the unusual total lack of male-male sharing as seen in Table 2. (see the levels of confidence in Fig 9).

**Fig 8.**
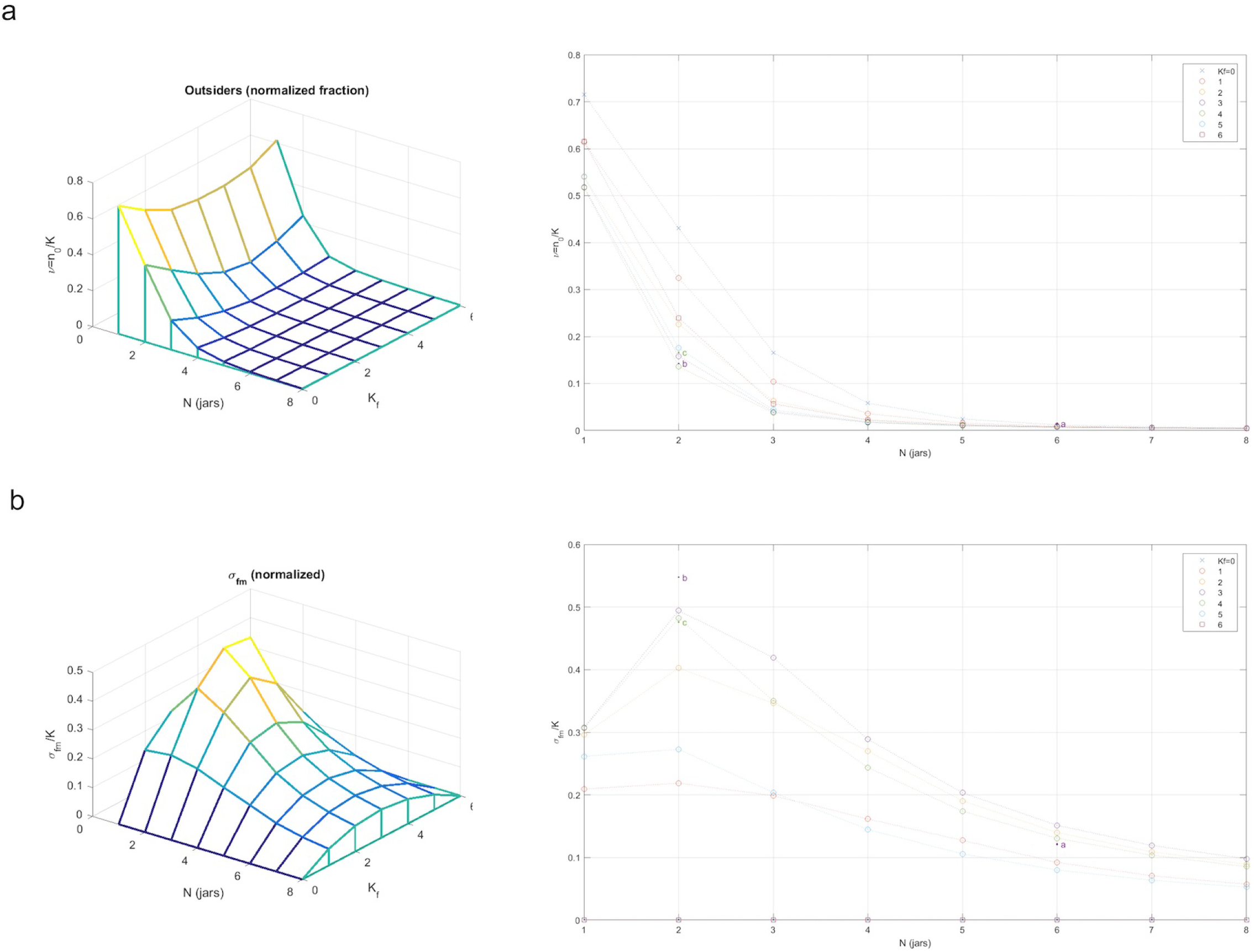
(a) **The average number of outsiders as a function of** (*N, K_f_*). 〈*n*_0_〉 is minimal for balanced sexs, *K_f_* = *K_m_*. The empirical points (*b, c*) are close to their corresponding calculated curves. (b) **The average female-male linkage** 〈*σ_fm_*〉 as a function of (*N, K_f_*). 〈*σ_fm_*〉 is maximal for balanced sexes *K_f_* = *K_m_* and *N* = 2. The empirical points (*b, c*) are again close to the calculated curve however, point (*a*) looks like an outliner.

Fig 9 demonstrates the tradeoff between gaining by having a high female-male linkage and losing due to a large number of outsiders. Thus, as one increases the density of animals, by reducing the number of pots, 〈*n*_0_〉 and 〈σ_*fm*_〉 start growing together and keep increasing monotonically, until reaching a turning-point (in our case, that point is specified as *N* = 2) where further increase of the density causes a decrease of 〈σ_*fm*_〉, accompanied by continuing increase of 〈*n*_0_〉. Fig 9 also presents the expected uncertainties in *n*_0_ and *σ_fm_* which are essential for making comparison with experiments. The uncertainty levels have two sources: one, due to the intrinsic fluctuations which occur as the animals keep moving between different occupancy configurations, and the other, due to errors in estimating the interaction parameters. Setting 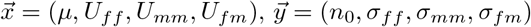, and expanding the error-matrix 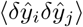 to the leading order in (1/*M*) one finds

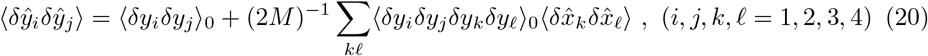

where 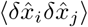 is the error-matrix in estimating the interaction-parameters, and 〈· · ·〉_0_ are the 2 and 4-point connected correlations for a set of known parameters. The last term on the RHS of (20) vanishes as the number of experiments *M* →∞. The first term, however, is controlled by the size of the system (decreases as *K, N* →∞, while the density *ρ* = *K/N* is kept finite) and, therefore, remains relatively large in many experimental designs. The case 1 ≤ *N* ≪ *K* is of particular interest. Referring to Eq 3 and setting *x* ≡ *μ/U*, we find that for weak interaction *ρ* = *x* + 1/2. However, as *U* increases (*U* ≃ 4*π*) the density crosses over to a staircase curve resembling the Mott-Hubbard transition [33] (Fig 10).

**Fig 9.**
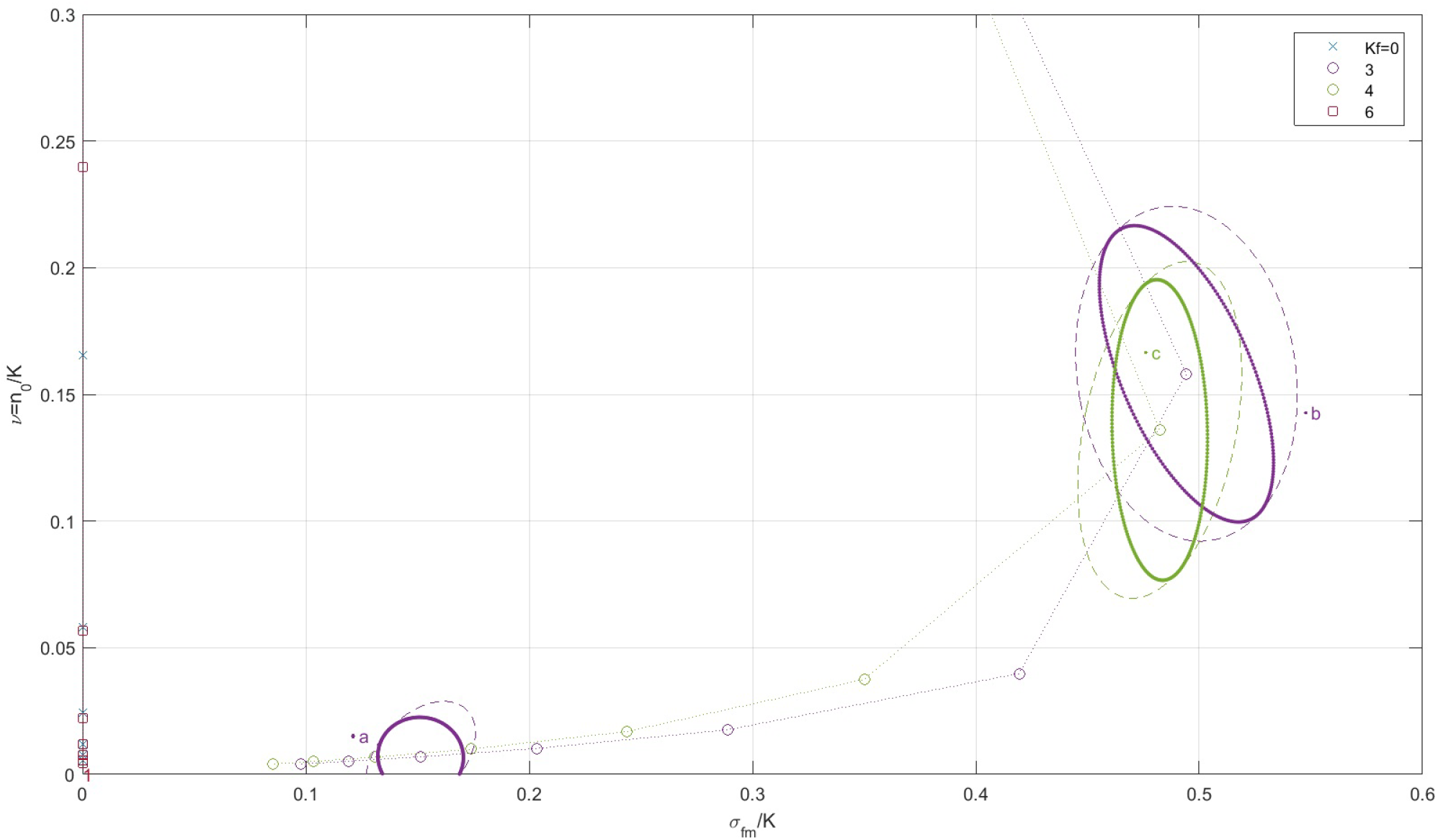
**The ‘equation-of-state’ in the 〈*σ_fm_*〉 – 〈*n*_0_〉 plane**, showing the tradeoff between high female-male linkage and large number of outsiders. The ellipses of 10% uncertainty demonstrate that empirical point (*b*) lays well within the range of error, whereas point (*a*) is a clear outliner (ellipse solid-line: finite-size 10% uncertainty for a given set of interaction-parameters, dashed-line: error in parameter estimation is included).

**Fig 10.**
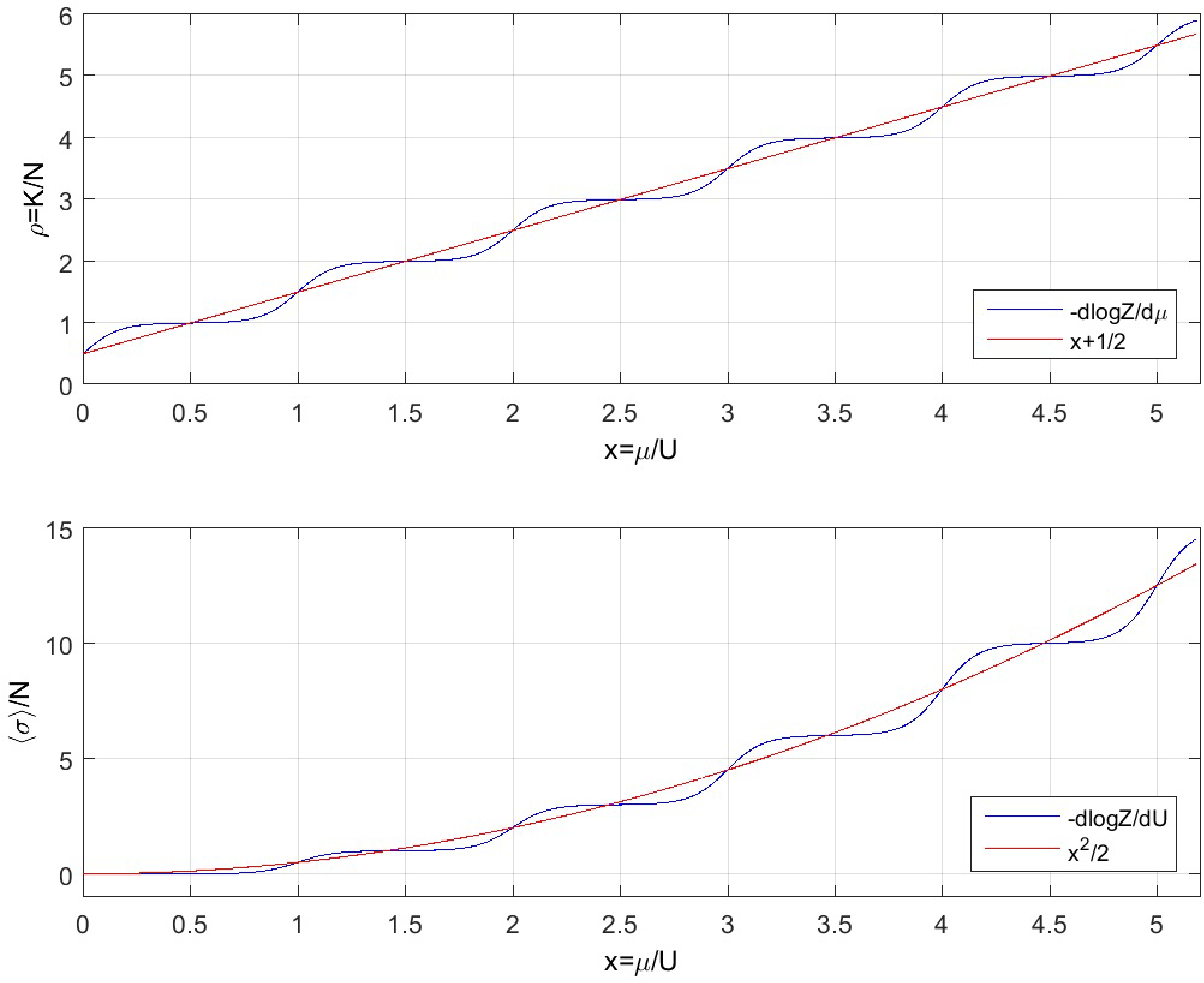
**The density** *ρ* = *K/N* **and the average linkage per den** = 〈*σ*〉/*N* **as a function of** *x* = *μ/U* **for large number of animals** (1 < *N* ≪ *K*). In red – weak interaction, *ρ* = *x* +1/2 and ξ = *x*^2^/2. In blue – strong repulsive interaction *U* ≃ 12.

## Conclusion

“Sociality” is an elusive concept; we [think we] know it when we see it. The difficulties that arise in trying to define it crystallize in the context of robotics, for example, wherein inanimate objects can exhibit collective swarmlike behaviors. Once living organisms are viewed as wetmare machinery, the arbitrariness inherent to any particular definition of “sociality” is uncontroversial. Nevertheless, once one has in mind a specific purpose, definitions of sociality customized to achieve clearly articulated predictions of behavior on explicitly stated terms may become possible.

Thus, as discussed at the 2018 Aspen Center for Physics workshop on “Physics of Behavior,” any quantitative measure of “sociality” is heavily dependent on context. We aim to develop reproducible laboratory measures that reflect (and eventually predict) field observations that could be relevant for successful commercial culture of the animal. The field observations reported here of octopus *O. laqueus* engaging in den-sharing, a behavior which is thought to be atypical of most octopus species, could indicate that they are more readily cultured in the lab without cannibalization, than are other species of octopus. Anecdotal evidence suggested that *O. laqueus* individuals tolerate one another: field observations of two animals apparently sharing the same den; the willingness of multiple individuals to cohabit for indefinitely within a single tank without a lid, a condition wherein many octopus species would – in our experience – flee the tank to certain death in a dark corner of the lab. The challenge is to move beyond anecdote. As with all biological systems, systematics often comes at the cost of artificial or unnatural settings. Octopuses that are not well-fed, for example, will generally try to eat one another in the lab, but EU guidelines forbid keeping octopus under conditions where eating one another is routine.

The anecdotal observation of den sharing in the field suggested to us that den sharing could be recast into a laboratory measure that might plausibly reflect certain aspects of sociality. In our hands, *O. laqueus* in laboratory tanks equipped with clay pots, exhibit distinctive behavior wherein they explore the dens in the evening hours before settling in for the night. Den sharing provides a readily measurable observable amenable to parameterization by number of dens and number of animals. To compensate for the apparent crudity of our measure, we were able to establish statistical uncertainty by assessing the *independence* of measurements with a suitably-defined *correlation time* without which statistical characterizations customary in the literature on sociality are rendered meaningless.

We studied the social tolerance of *O. laqueus* by measuring the den occupancy of dens in the lab for varying densities of animals and several sex-mixtures. We found that *O. laqueus* tolerate other individuals by sharing tanks and dens, with typically no loss to cannibalism or escape. However, animals also exhibit significant levels of social repulsion, and individuals often chose a solitary den when given the option. The patterns of den occupancy are shown to follow a model of maximum entropy. The animals were modeled as ‘identical particles’ with on-site pair interaction. The three interaction-parameters, that determine the amount of social attraction/repulsion between animals according to the sex, together with the chemical potential, that keeps the animals from leaving the dens – were estimated from the experiment by a standard maximum likelihood calculation. The parameters obtained in this way were then used to characterize the social behavior in large set of experimental conditions and to identify potential outliers. This procedure, as well as the general applicability of a maximum entropy model in this context, remain to be verified in future experiments with larger sample statistics.

## Supporting information

**S1 Video. Two *O. laqueus* in close proximity in the field, possibly sharing a den.**

**S2 Video Elastomere injection of O. *laqueus.***

## Acknowledgments

We wish to thank J. Simmons, D. Calzarette, J. Gordon, and C. Timmons, for assistance in animal care and experiments, and G. Ilsley for initial work on data modeling.

NO and RY made initial field and lab observations of social tolerance in O.laqueus. EE and RY made additional field observations and collected animals. EE, NO, RY and KD designed and EE and KD performed all lab experiments. EE performed initial analysis and RP and JM performed detailed analysis and modeling. EE, RP, and JM all contributed to writing the manuscript.

